# Raf-MEK-ERK signaling mediates FLT3-induced mechanical allodynia and reduced morphine analgesic efficacy in male rodents

**DOI:** 10.64898/2026.10.04.756546

**Authors:** Jean-Philippe Leyris, Danaë Ajas, Juliette Bertin, Chamroen Sar, Sylvie Mallié, Thiago M. Cunha, Fabrice Ango, Hamid Moha ou Maati, Ilana Méchaly, Jean Valmier, Cyril Rivat

## Abstract

The cytokine FL and its receptor, Fms-like tyrosine kinase 3 (FLT3), contribute to chronic pain and to the maladaptive effects of morphine but the intracellular mechanisms underlying these effects remain incompletely understood. Here, we investigated the contribution of the Raf-MEK-ERK pathway to FLT3dependent nociceptive signaling and to the modulation of morphine analgesia. Using HEK293T cells expressing human or mouse FLT3, primary dorsal root ganglion (DRG) neurons, and rodent models, we examined ERK activation, TRPV1 sensitization, and behavioral responses following FLT3 activation by its ligand, FL. FL induced concentration-dependent ERK phosphorylation through FLT3, while pharmacological inhibition of FLT3, Raf, or MEK reduced this response. In primary DRG neurons, MEK inhibition attenuated FLinduced potentiation of capsaicin-evoked calcium responses, supporting a role for MEK-ERK signaling in TRPV1 sensitization. In vivo, intrathecal FL increased ERK phosphorylation in DRGs and induced mechanical hypersensitivity, both of which were reduced by Raf or MEK inhibition. In FLT3/MOR-expressing cells, morphine potentiated FL-induced ERK activation, an effect reduced by naloxone and abolished by FLT3 inhibition. In rats, MEK inhibition prevented the FL-induced reduction of morphine analgesia and subsequent mechanical hypersensitivity. Together, these findings identify Raf-MEK-ERK signaling as an important downstream pathway linking neuronal FLT3 activation to nociceptor sensitization and impaired morphine analgesic efficacy.

**Highlights:**

- Mouse or human FLT3 stimulation activates ERK which is reversed by FLT3, Raf or MEK inhibition
- FL increases capsaicin-induced Ca2+ influx via MEK-ERK signaling in DRG neurons
- FL injection activates ERK in DRG for at least 1h
- FL-induced pain hypersensitivity is partially prevented by Raf or MEK inhibition
- FL reduces morphine analgesia which is totally prevented by MEK inhibition

## Introduction

FLT3 (Fms-related tyrosine kinase 3), also known as FLK-2 or STK-1, is a class III receptor tyrosine kinase (RTK). It is primarily expressed in haematopoietic stem cells in the bone marrow, as well as in various haematopoietic organs, such as the spleen and thymus. FLT3 is specifically activated by its endogenous ligand, FL (also known as FLT3L) that is a cytokine expressed mainly by immune cells. The FL-FLT3 signaling is a major regulator of hematopoietic and immune homeostasis, controlling the proliferation, survival, and differentiation of hematopoietic progenitor cells, especially in the myeloid and B-cell compartments^1–4^. In the nervous system, FLT3 is expressed in both central and peripheral structures^5–9^ and plays a critical role in neural development, notably by promoting the survival of neuronal progenitors^10^. While these studies highlight its developmental functions, increasing evidence suggests that FLT3 signaling may also regulate neuronal activity in the adult nervous system. In line with this, our previous work demonstrated that FLT3 signaling contributes to the modulation of nociceptive pathways^6,7,9^

We have showed FLT3 to be expressed in sensory neurons of the dorsal root ganglia (DRG), playing a critical role in the regulation of nociception and in chronic pain. Specific activation of FLT3 signaling in primary sensory neurons of DRG enhances neuronal excitability, thereby promoting pain hypersensitivity. Conversely, blocking FLT3 expression or its activity alleviated pain hypersensitivity in mouse model of peripheral neuropathic pain^6^ as well as in the surgical pain model^7^. Recently, we also demonstrated the contribution of FLT3 in the development of morphine-induced analgesic tolerance and sustained hyperalgesia^8^. Through a functional interaction with µ opioid receptor (MOR) expressed in the sensory neurons of the DRG, FLT3 promotes morphine-induced neuronal hyperexcitability via cAMP-dependent mechanisms. Interestingly, pharmacological blockade of FLT3 not only blunts morphine tolerance and hyperalgesia but also enhances morphine analgesia without affecting the other morphine side effects such as the motivational effects, withdrawal somatic symptoms and respiratory depression. However, the specific contribution of FLT3 signaling pathways in the modulation of nociception and morphine analgesic effects remain to be described.

FLT3 exerts its cellular effects through three major intracellular signaling pathways: PI3K/AKT/mTOR, JAK/STAT5, and MAPK/ERK1/2, which collectively regulate its physiological and pathological roles in haematopoiesis. Beside its role in chronic pain^11–13^, ERK pathway has also been involved in opioid effects. Chronic morphine administration produces ERK activation in brain structures such as locus coeruleus and caudate/putamen, periaqueductal gray^14^ promoting withdrawal and tolerance to morphine analgesia. In sensory neurons of the DRG, ERK is activated in DRG cultures after repeated morphine exposure in CGRP positive neurons^15^. The inhibition of ERK totally reduces up-regulated expression of CGRP and activation of the transcription factor CREB after chronic morphine administration and its inhibition reduces morphine tolerance^16^. In addition, the critical role of ERK in morphine analgesic tolerance has also been linked to altered expression of TRPV1 in DRG, spinal cord and sciatic nerve^17^. Blocking ERK activation not only reduced morphine tolerance but also prevented morphine-induced TRPV1 overexpression. Together, these findings suggest that ERK appears to be a key intracellular mediator of morphine effects in sensory neurons of the DRG.

Building on our previous evidence that FLT3 modulates opioid analgesia^8^, this study aimed to determine whether activation of the Raf-MEK-ERK pathway functionally links FLT3 signaling to nociceptive hypersensitivity and reduced morphine analgesic efficacy.

## Material and Methods

### Animals

Experiments were performed in male Sprague-Dawley rats (150-170g, Janvier, France) or male C57BL/6 naive mice (25-30g, Janvier, France). All the procedures were approved by the French Ministry of Research APAFIS#20114-2019040310181741 v3. Animals were maintained in a climate-controlled room on a 12 h light/dark cycle and allowed access to food and water ad libitum.

### Drugs

Human or mouse recombinant FL (FLT3 ligand) were obtained from R&D. Human recombinant FL was also produced in the E. coli Rosetta (DE3) strain (Novagen) in our laboratory using the pET15b-rhFL plasmid according to the protocol described (Verstraete et al., 2009) with some minor modifications. The rh-FL was checked for endotoxin content using the Pyrogen Recombinant Factor C endotoxin detection assay from LONZA (Walkersville MD, USA) and was found free of endotoxins. For intrathecal injection, the rh-FL (50 ng) was dissolved in saline and a 30□G needle attached to a Hamilton syringe was inserted between vertebrae in lightly restrained, isoflurane-anaesthesized mice or rats. The reflexive tail flick was used to confirm the punction. A total volume of 10 (mice) or 20□ µl (rat) was injected. Morphine sulfate was purchased from Francopia, dissolved in 0.9% sodium chloride and administered subcutaneously. For acute morphine-FL treatment protocol in rats, one subcutaneous morphine injection (2 mg/kg) was performed 30 minutes after FL intrathecal injection. The MEK inhibitor PD98059 (#P215 - Sigma-Aldrich) was administered at a concentration of 10µg/20µl, the Raf inhibitor TAK632 (ci #S7291 - Selleck Chem) was administered at a concentration of 500 ng/10µl. BDT001 was synthesized by D. Rognan at Strasbourg (Laboratoire d’Innovation Thérapeutique, UMR7200, CNRS - Université de Strasbourg, Illkirch 67400 France). BDT001 was dissolved in 7% Tween 20 in a saline solution.

### Behavioral analysis

Before testing, rats and mice were acclimatized for 60 min in the temperature and light-controlled testing room within a plastic cylinder or on wire mesh. Experimenters were blinded to the drug administered.

#### Tactile sensitivity

Tactile withdrawal threshold was determined in response to probing of the hindpaw with eight calibrated von Frey filaments (Stoeling, Wood Dale, IL, USA) in logarithmically spaced increments ranging from 0.41 to 6 g (4-150 mN). Filaments were applied perpendicularly to the plantar surface of the paw. The 50% paw withdrawal threshold was determined in grams by the Dixon nonparametric test26. The protocol was repeated until three changes in behavior occurred. This method was used both in rats and mice with adapted filaments.

#### Paw pressure test

To assess mechanical sensitivity in rats, a variant method of the Randall-Selitto method was performed as previously described^18^. Briefly, increasing pressure was applied to the hind paw until the rat squeaks. The Basile analgesimeter (Apelex, Massy, France; stylus tip diameter, 1 mm) was used. A 600-g cutoff value was determined to prevent tissue damage. After their arrival to the laboratory, rats were kept in their colony room for 4 days. To avoid any experimental stress, experimentations were realized by the same experimenter in quiet conditions. Animals were habituated to both experimenters, experimental room and devices for at least 10 days before starting the experiments. Before starting the experiments, basal nociceptive threshold was measured during 2 days in a row and on the experimental day to verify nociceptive threshold stability. To assess morphine analgesia, nociceptive threshold was assessed every 30 minutes for 2h30 until return to a basal nociceptive threshold. Basal nociceptive threshold was assessed once daily after FL-morphine administration.

### HEK293T cell culture

HEK293T cells, used for Phospho-ERK assays, were maintained in DMEM Glutamax (Invitrogen) supplemented with antibiotics (penicillin 50 U/ml, streptomycin 50 μg/ml) and 10% heat-inactivated Foetal Calf Serum. The HEK 293 T cells were transiently transfected with human FLT3 receptor cDNA and human MOR receptor cDNA (see below) using Lipofectamine 2000 (Invitrogen) according to manufacturer’s instructions. Transfections were performed in 96-well plates using a cell density of 50,000 cells per well. Prior to cell plating, wells were pre-coated with Poly-L-Ornithine (50 μl of 10 mg/ml) for 30 min at 37°C.

### Transfection procedures

Transfection mixes were prepared using 50 ng of SNAP-FLT3 plasmid, 50ng of MOR-YFP, 0.25 μl of Lipofectamine 2000 (Invitrogen) and 50 μl Opti-MEM per well. Prior to their addition to plates, transfection mixes were preincubated for 20 min at room temperature. Then 100 μl of HEK293T cells at a density of 500,000 cells/ml were plated in each well and were incubated at 37°C under 5% CO2 for 24h.

### ERK1/2 time-resolved FRET assays

To determine the degree of phosphorylation of ERK after the stimulation of cells expressing MOR and FLT3 receptors, by different compounds, we used the CisBio assay (Phospho-ERK1/2 (Thr202/Tyr 204) #64ERKPEH). Cells transiently transfected with the different receptors were cultured in a 96-well plates. Twenty-four hours after cell transfection, cells were starved overnight in DMEM-Glutamax serum free medium and then incubated in a total volume of 100□μl/well of the DMEM-Glutamax serum free medium, containing or not the different pharmacological compounds at the different concentrations as indicated. More precisely, cells were incubated with 10 µM of naloxone (#N7758 - Sigma-Aldrich), 10 µM of PD98059 (#P215 - Sigma-Aldrich ; MEK inhibitor), 0.1 µM of HS-173 (PI3Kinase inhibitor; #S7356 - Selleckchem), 10 µM of AG490 (658411 - Calbiochem), 50 µM of BDT001 (synthesized by Laboratoire d’Innovation Therapeutique, Strasbourg) and 1 µM of AC220 (#S1526 - Selleckchem, FLT3 inhibitor) for 90min at 37°C, prior to addition of 10 µM of morphine during 15 min at 37°C, followed by 5nM of rhFL during 10min at 37°C. Cells were incubated at 37°C before the stimulation mix was removed and cells were then lysed in 50□μl/well of the supplemented lysis buffer for 60min at room temperature with shaking. Then 16□ μl of each lysate was transferred into a 384-well small volume white plate and made up to the final volume of 20□μl/well with 2□μl of anti-ERK1/2-Europium/Terbium Cryptate and 2□μl of anti-Phospho-ERK1/2-d2 antibody solutions prepared in the detection buffer. The plate was then incubated for 2□h at room temperature before reading the fluorescence emission at 620 and 665□nm using a PHERAstar FS plate reader (BMG Labtech). Data are presented as the percentage of Non Stimulated from HTRF ratios, which are calculated as the ratio between the emission at 665□nm and the emission at 620□nm (×10,000).

### Adult sensory neuron culture

For calcium imaging, neuron cultures were established from lumbar (L4–L6) DRG^6^. Briefly, ganglia were successively treated by two incubations with collagenase A (1□mg/ml, Roche Diagnostic, France) for 45□ min each (37□°C) and trypsin-EDTA (0.25%, Sigma, St Quentin Fallavier, France) for 30□min. They were mechanically dissociated through the tip of a fire-polished Pasteur pipette in neurobasal (Life Technologies, Cergy-Pontoise, France) culture medium supplemented with 10% fetal bovine serum and DNase (50□ U/ml, Sigma). Isolated cells were collected by centrifugation and suspended in neurobasal culture medium supplemented with 2% B27 (Life Technologies), 2□mM glutamine, penicillin/streptomycin (20 U/ml, 0.2□mg/ml) plated at a density of 2500 neurons per coverslip and were incubated in a humidified 95% air 5% CO2 atmosphere at 37□°C. The data were obtained from experiments conducted on 12 mice.

### Calcium imaging

For calcium imaging video microscopy [Ca^2+^]_i_ fluorescence imaging, DRG neurons were loaded with fluorescent dye 2.5□µM Fura-2 AM (Invitrogen, Carlsbad, CA) for 30□min at 37□°C in standard external solution contained: 145□ mM NaCl, 5□ mM KCl, 2□mM CaCl2, 2□mM MgCl2, 10□mM HEPES, 10□mM glucose (pH adjusted to 7.4 with NaOH and osmolarity between 300 and 310□mOsm). The coverslips were placed on a stage of Zeiss Axiovert 200 inverted microscope (Zeiss, München). Observations were made at room temperature (20-23□°C) with a ×20 UApo/340 objective. Fluorescence intensity at 505□nm with excitation at 340 and 380□ nm were captured as digital images (sampling rates of 0.1-2□s). Regions of interest were identified within the soma from which quantitative measurements were made by re-analysis of stored image sequences using MetaFluor Ratio Imaging software. [Ca^2+^]_i_ was determined by ratiometric method of Fura-2 fluorescence from calibration of series of buffered Ca^2+^ standards. Neurons were distinguished from non-neuronal cells by applying 25□mM KCl, which induced a rapid increase of [Ca^2+^]_i_ only in neurons. Pulses of capsaicin were applied at 2□min-intervals and capsaicin and drugs was added after the third pulse. The same temporal stimulation protocol was used for TRPV1 (capsaicin). All drugs and solutions were applied with a gravity-driven perfusion system. For data analysis, amplitudes of [Ca^2+^]_i_ increases, ΔF/Fmax, caused by stimulation of neurons with capsaicin were measured by subtracting the “baseline” F/Fmax (mean for 30□s prior to capsaicin addition) from the peak F/Fmax achieved on exposure to capsaicin. In the absence of any treatment the distribution of these ratios was well fitted by a normal distribution.

### Western Blot

ERK phosphorylation in mouse DRGs was assessed by Western blotting using two independent experimental series. C57BL/6 mice received an intrathecal injection of recombinant FL (50 ng; R&D Systems, #308-FK) or saline in a final volume of 5 µL, and bilateral lumbar DRGs (L4-L6) were collected 30 min or 1 h after injection, respectively. Tissues were homogenized with ceramic beads in RIPA buffer supplemented with phosphatase and protease inhibitors, using either a MagNA Lyser Instrument (Roche; two 30-s cycles at 6,000 rpm; Lysing Matrix D, MP Biomedicals, #116913100) or a tissue lyser (two 8-min cycles at 25 s□¹). Lysates were clarified by centrifugation at either 10,000 × g for 15 min or 500 × g for 5 min at 4°C, and protein concentrations in the supernatants were determined using a bicinchoninic acid assay. Proteins were separated by SDS-PAGE, transferred onto nitrocellulose membranes, and blocked with 5% bovine serum albumin in TBS containing 0.1% Tween-20 for 1–2 h at room temperature. Membranes were incubated overnight at 4°C with antibodies against phospho-p44/42 MAPK (ERK1/2; Thr202/Tyr204), total p44/42 MAPK (ERK1/2), and either α-tubulin or βIII-tubulin. After washing, membranes were incubated with the appropriate fluorescent IRDye-conjugated secondary antibodies for 2 h or horseradish peroxidase-conjugated secondary antibodies for 1 h at room temperature. Signals were detected using either an Odyssey CLx Imaging System (LI-COR Biosciences) or ECL reagents and a ChemiDoc XRS system (Bio-Rad), and quantified using Image Studio or Image Lab software, respectively. Phosphorylated ERK levels were normalized to total ERK, whereas total ERK levels were normalized to the corresponding tubulin loading control.

### Use of generative artificial intelligence

ChatGPT (OpenAI) was used to assist with code generation for sequence alignment, structural comparison and figure preparation. The analyses were subsequently performed using the computational methods and software described above. All code, numerical results and graphical outputs were reviewed and validated by the authors, who take full responsibility for their accuracy and interpretation.

### Sequence retrieval and pairwise alignment

The canonical human and mouse FLT3 protein sequences were retrieved from UniProtKB/Swiss-Prot (human FLT3, accession P36888, 993 amino acids; mouse Flt3, accession Q00342, 1,000 amino acids). Global pairwise sequence alignment was performed in Biopython v1.88 using the PairwiseAligner implementation of the Needleman–Wunsch algorithm, the BLOSUM62 substitution matrix, a gap-opening penalty of 10, and a gap-extension penalty of 0.5. Sequence identity was calculated as the number of identical aligned positions divided by the total alignment length, including gap-containing positions. Conservative substitutions were defined as aligned residue pairs with a positive BLOSUM62 substitution score. The protein kinase domains were defined according to UniProt annotations (human FLT3, residues 610-943; mouse Flt3, residues 611-946) and aligned separately using the same parameters. The resulting sequence identities were 85.5% for the full-length receptors and 92.6% for the kinase domains.

### AlphaFold model retrieval and structural comparison

Full-length AlphaFold models of human and mouse FLT3 were downloaded from the AlphaFold Protein Structure Database (human, AF-P36888-F1-model_v6; mouse, AF-Q00342-F1-model_v6). Structural comparison was restricted to the structured kinase core to avoid the influence of a flexible kinase-domain insertion. The residues included in the structural alignment were 610-710 and 762-943 for human FLT3 and 611-711 and 763-946 for mouse Flt3. The flexible intervening regions, corresponding to residues 711-761 in human FLT3 and 712-762 in mouse Flt3, were excluded from the RMSD calculation. Residue correspondence was defined from the pairwise sequence alignment, and the mouse kinase core was superposed onto the human kinase core in PyMOL v3.2.0 by least-squares fitting of corresponding Cα atoms. The final structural comparison included 283 aligned Cα pairs and yielded an RMSD of 1.28 Å. Structural representations were generated in cartoon mode, with human FLT3 shown in gray and mouse Flt3 shown in green.

### Statistical analysis

All experiments were randomized. Data are expressed as the mean□±□ s.e.m. All sample sizes were chosen based on our previous studies except for animal studies for which sample size has been estimated via a power analysis using the G-power software. The power of all target values was 80% with an alpha level of 0.05 to detect a difference of 50%. Statistical significance was determined by analysis of variance (ANOVA one-way or two-way for repeated measures, over time), followed by Bonferonni’s post-hoc test to compare with the control group.

## Results

### FL-induced ERK activation in HEK293T cells expressing human or mouse FLT3

To assess the cross-species conservation of FLT3, we compared the amino acid sequences and AlphaFold-predicted structures of the human and mouse receptors. Sequence alignment revealed a high degree of conservation, with 85.5% identity across the full-length proteins and 92.6% identity within their kinase domains (Figure 1A). Consistent with this sequence conservation, the predicted kinase domains displayed highly similar three-dimensional folds. Structural comparison was restricted to the structured kinase core, excluding the flexible kinase-domain insertion. Superposition of the human and mouse kinase cores included 283 aligned Cα pairs and yielded an RMSD of 1.28 Å, indicating strong structural conservation (Figure 1B). We next investigated the ability of FL to activate ERK signaling in HEK293T cells expressing either human (Figure 1C) or mouse (Figure 1D) FLT3. Commercially available recombinant human and mouse FL were compared with internally produced recombinant human FL. All three FL preparations induced a concentration-dependent increase in ERK activation in cells expressing either receptor ortholog. Comparable response profiles were observed with human and mouse FLT3, irrespective of the species origin of the ligand (Figure 1C, D). Together, these findings demonstrate strong structural and functional conservation of the FL-FLT3 signaling axis between humans and mice. They also show that human and mouse FL can activate FLT3 across species^19,20^, supporting the use of mouse experimental systems and internally produced human recombinant FL in subsequent experiments.

**Figure 1.**
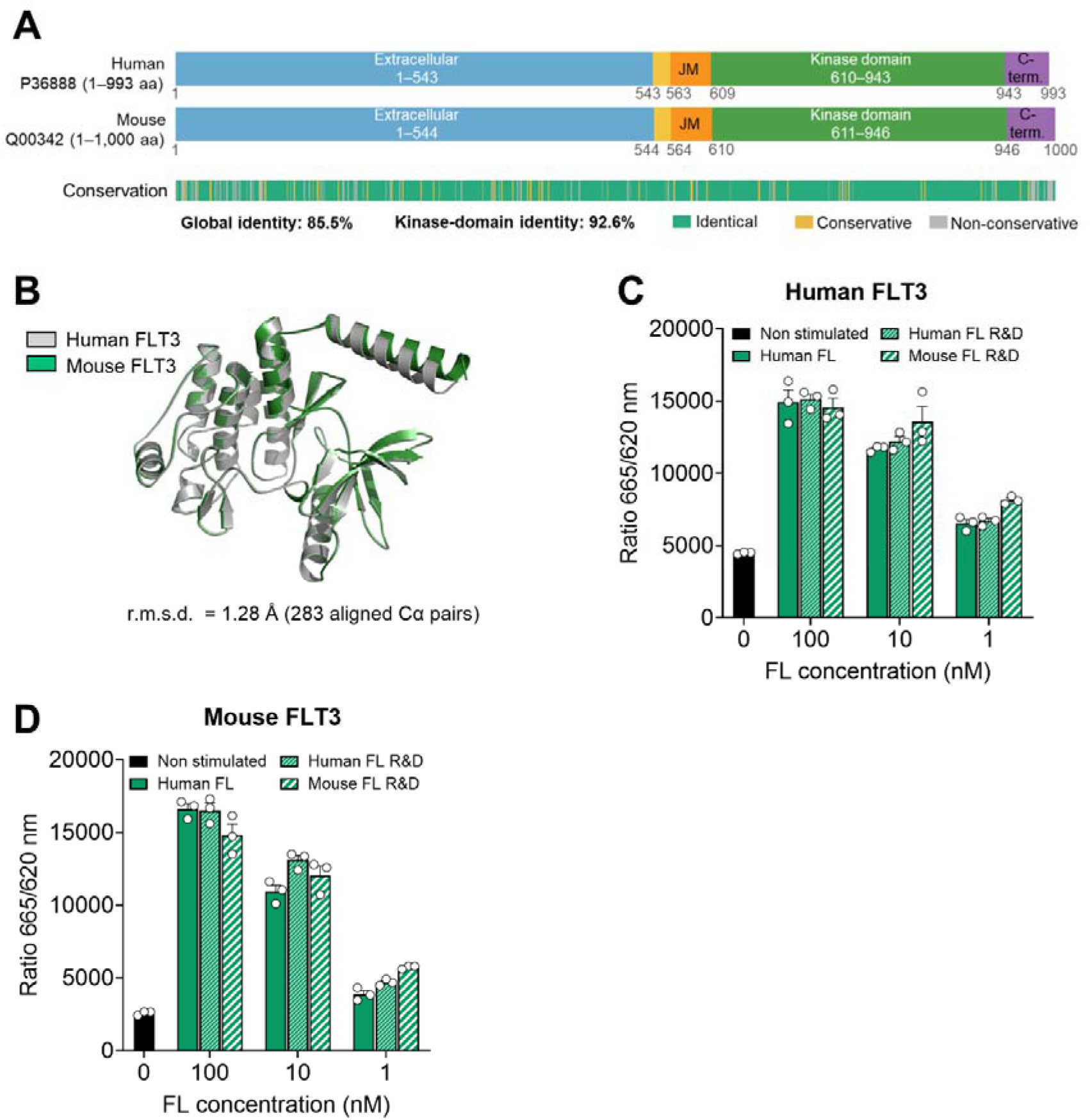
Structural and functional conservation of FLT3 signaling between humans and mice. **(A)** Schematic comparison of the canonical human and mouse FLT3 protein sequences, showing the extracellular, transmembrane, juxtamembrane, kinase, and C-terminal regions. Aligned residues are classified as identical, conservative substitutions, or non-conservative substitutions. Human and mouse FLT3 display 85.5% global sequence identity and 92.6% identity within the kinase domain. Amino acid boundaries are indicated below each receptor schematic. **(B)** Structural superposition of the AlphaFold-predicted kinase cores of human FLT3 (gray; AF-P36888-F1, v6) and mouse Flt3 (green; AF-Q00342-F1, v6). The UniProt-annotated kinase domains comprise residues 610-943 in human FLT3 and 611-946 in mouse Flt3. The structural comparison was restricted to the structured kinase core, corresponding to residues 610-710 and 762-943 in human FLT3 and residues 611-711 and 763-946 in mouse Flt3, thereby excluding the flexible kinase-domain insertion. Superposition of 283 aligned Cα pairs yielded an RMSD of 1.28 Å. **(C, D)** Activation of human and mouse FLT3 by human and mouse FL. HEK293T cells expressing human FLT3 (**C**) or mouse Flt3 (**D**) were left unstimulated or stimulated with 1, 10, or 100 nM internally produced recombinant human FL, commercially available recombinant human FL (R&D Systems), or commercially available recombinant mouse FL (R&D Systems). FLT3-dependent ERK activation was assessed using an HTRF phospho-ERK1/2 assay and expressed as the fluorescence ratio at 665/620 nm. Data were obtained from one experiment. Individual technical replicates are shown as open circles, and bars represent mean ± SEM.

To confirm that FL-induced ERK phosphorylation is due to the activation of FLT3 receptor, we assessed the effects of the FLT3 inhibitor BDT001 on ERK phosphorylation. BDT001 markedly reduced the phosphorylation of ERK induced by FL with an estimated IC□□ of 7.8 ± 3.7 µM (n = 2 biological replicate). Then, to determine whether Raf contributes to FLT3-dependent ERK signaling, we examined the effect of pharmacological inhibition of this pathway, previously reported to be activated downstream of FLT3^21^ (Figure 2A). In FLT3-expressing HEK293T cells, the pan-Raf inhibitor TAK-632 dose-dependently decreased FL-induced ERK phosphorylation with an IC50 of 0.8 ± 0.2 µM. Similarly, the application of MEK inhibitor P98059 inhibited FL-induced ERK phosphorylation with an IC50 6.5 ± 2.20 µM. These results identify Raf as a signaling intermediate linking FLT3 activation to ERK phosphorylation and support the engagement of the Raf-MEK-ERK pathway downstream of FL/FLT3 signaling.

**Figure 2.**
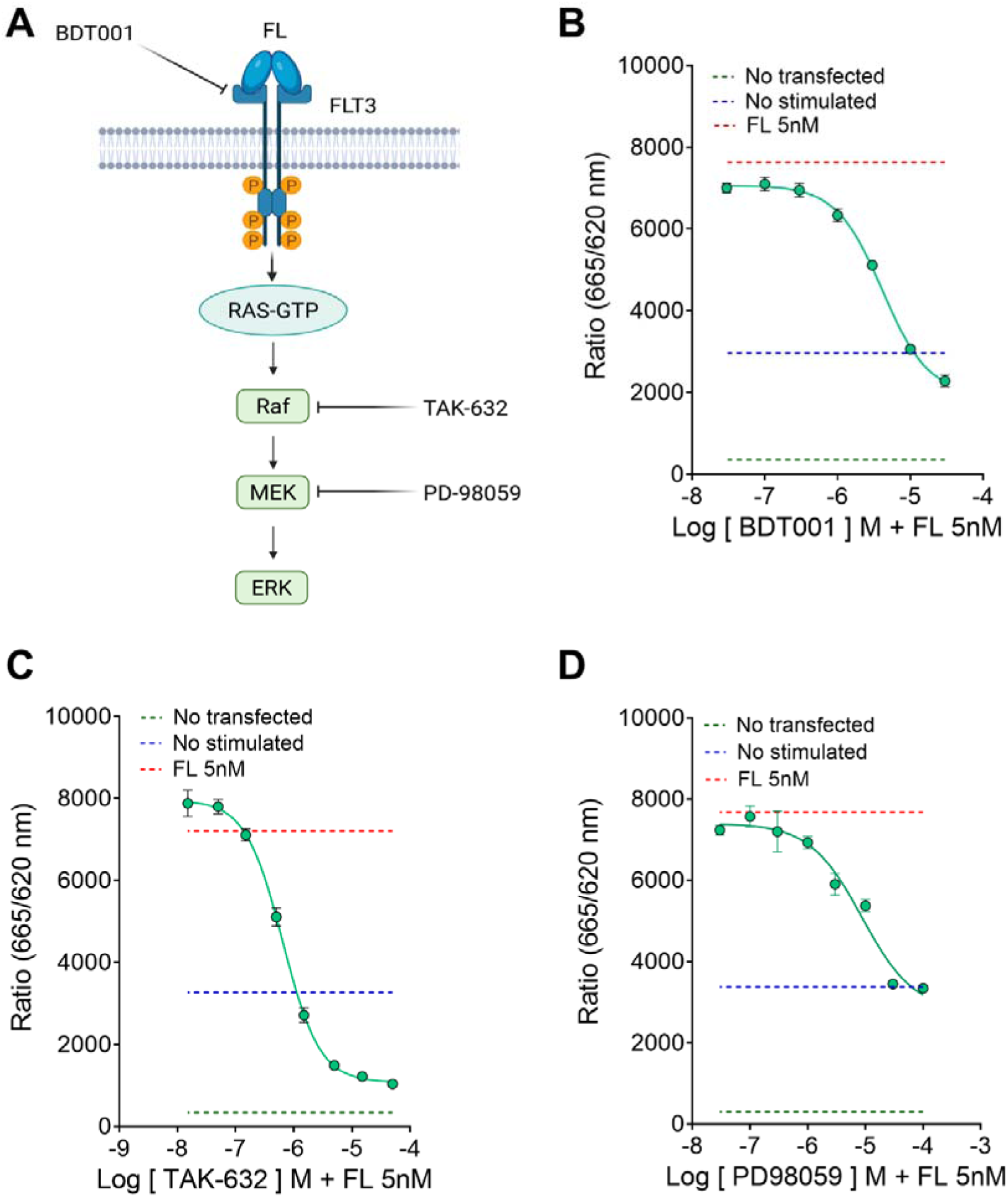
FL induces ERK activation in HEK293T cells through the FLT3–Raf–MEK signaling pathway. (**A**) Schematic representation of the FLT3-Ras-Raf-MEK-ERK signaling pathway and the pharmacological targets of the pan-Raf inhibitor TAK-632 and the MEK inhibitor PD98059. (**B-D**) Concentration-dependent inhibition of FL-induced ERK activation in HEK293T cells expressing human FLT3. Cells were stimulated with FL (5nM) in the presence of increasing concentrations of the FLT3 inhibitor BDT001 (B), TAK-632 (C) or PD98059 (D). ERK activation was measured using an HTRF phospho-ERK1/2 assay and expressed as the fluorescence ratio at 665/620 nm. Dashed lines indicate the responses measured in non-stimulated cells (NS), non-treated cells (NT), and cells stimulated with FL alone, as indicated. Curves were fitted using a nonlinear regression model. Data are presented as mean ± SEM from one representative experiment out of two independent experiments.

### ERK signaling contributes to FL-induced potentiation of TRPV1 function in mouse DRG sensory neurons

We previously demonstrated that FL concentration-dependently potentiates TRPV1-mediated intracellular calcium ([Ca^2+^]_i_) responses to repeated capsaicin stimulation^6^. Because TRPV1 function can be modulated by the MEK-ERK signaling pathway^22,23^, we investigated whether this pathway contributes to FL-induced TRPV1 sensitization. Calcium imaging experiments were performed in primary cultures of adult mouse DRG neurons treated with PD-98059, an inhibitor of MEK-dependent ERK activation. PD-98059 alone did not alter the [Ca^2+^]i response evoked by capsaicin (Figure 3A). In contrast, inhibition of MEK markedly reduced the potentiation of capsaicin-evoked [Ca^2+^]i responses induced by recombinant FL (Figure 3B). These findings indicate that activation of the MEK-ERK pathway contributes, at least in part, to FL-induced sensitization of TRPV1 in sensory neurons.

**Figure 3.**
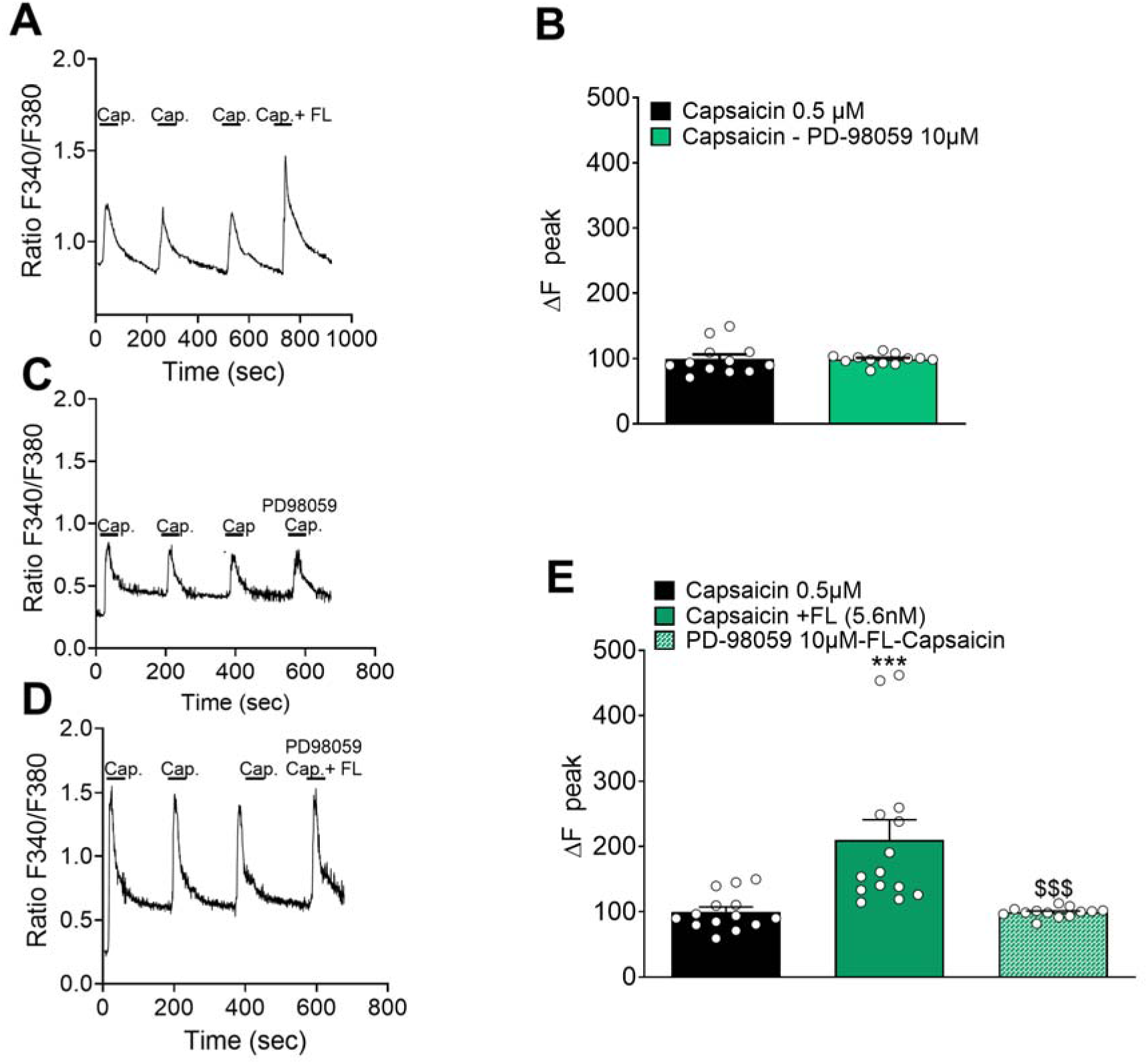
Capsaicin-induced intracellular calcium dynamics in primary cultures of DRG sensory neurons. (**A**) Traces of [Ca^2+^]_i_ responses to repeated bath applications of capsaicin following by a fourth peak corresponding to the Capsaicin + PD-98059 perfusion. (**B**) Quantification of F340/F380 fluorescence ratio calcium response measured at the fourth application of Capsaicin 0.5 µM (control condition, n=12 cells) or Capsaicin 0.5 µM + PD98059 10 µM (n=12 cells) as in A. Results are expressed as response amplitudes normalized to capsaicin alone (ΔF peak). (**C-D**) Traces of [Ca^2+^]_i_ responses to repeated bath applications of capsaicin 0.5 µM following by a fourth peak corresponding to the Capsaicin + FL perfusion (C) or Capsaicin + FL + PD-98059 (D). (**E**) Quantification of F340/F380 fluorescence ratio calcium response measured at the fourth application of Capsaicin 0.5 µM (control condition, n=14 cells), Capsaicin 0.5 µM + FL 5,6 nM (n=12 cells) and Capsaicin 0.5 µM + FL 5,6 nM + PD-98059 10 µM (n=13 cells) as in C and D. Results are expressed as response amplitudes normalized to capsaicin alone (ΔF peak). Number of experiments n = 12primary culture of DRG neurons, statistical analysis: one-way ANOVA followed by Bonferroni test for multiple comparisons, *** P < 0.001; $$$ P < 0.001. Cap.: Capsaicin

### FL induces mechanical hypersensitivity through activation of the Raf-MEK-ERK pathway in mouse DRGs

We next sought to determine whether FLT3 stimulation activates ERK signaling in vivo. Therefore, we assessed ERK phosphorylation in the DRGs by Western Blot analysis following intrathecal administration of FL. The phospho-ERK/total ERK ratio was significantly increased at 30 and 60 min after FL administration compared with saline-treated controls (Figure 4A). Of note, there was no difference between groups regarding the ratio total ERK on tubulin suggesting no change in the expression of ERK (data not shown). Next, we investigated the effects of the Raf-MEK-ERK pathway in FL-induced mechanical pain hypersensitivity in naïve male mouse (Figure 4B). Consistent with our previous findings^6,7,9^, intrathecal FL administration induced robust mechanical hypersensitivity 4h and 24h after its administration, as evidenced by an increased percentage of withdrawal responses to stimulation with a 0.6-g von Frey filament. Pharmacological inhibition of either Raf with TAK-632 or MEK with PD-98059 significantly attenuated FL-induced mechanical hypersensitivity 2h after FL administration (Figure 4C). Together, these findings indicate that intrathecal FL activates ERK signaling in the DRGs via the engagement of the Raf-MEK-pathway contributing to FL-induced mechanical hypersensitivity *in vivo*.

**Figure 4:**
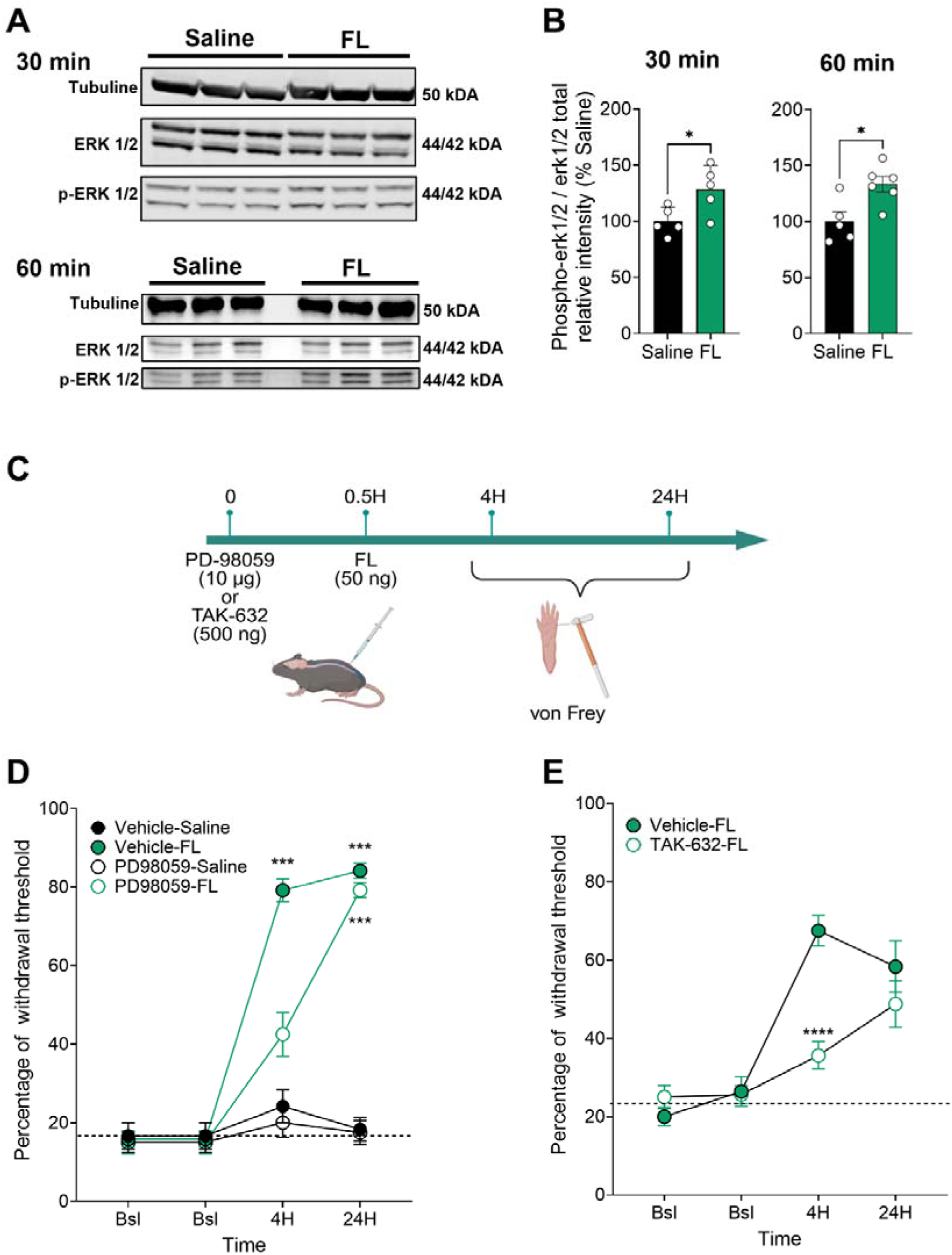
FL-induced Raf-MEK-ERK signaling pathway activation promoted mechanical pain hypersensitivity. (**A**) Representative Western blot bands showing ERK, p-ERK1/2 and tubulin (loading control) protein levels in L4-L6 DRG 30 and 60 min after intrathecal FL administration. (**B**) Densitometric quantification of the p-ERK/total ERK ratio, expressed as a percentage of the Saline group. Statistical analysis was performed using unpaired t-test *P < 0.05. (**C**) Experimental timeline used to evaluate the contribution of Raf-MEK-ERK signaling to FL-induced mechanical hypersensitivity. PD98059 (10 µg) or TAK-632 (500 ng) was administered 30 min before FL (50 ng). Mechanical withdrawal thresholds were assessed using von Frey filaments before treatment and at 4 and 24 h after FL administration. (**D**) Effect of the MEK inhibitor PD98059 on FL-induced mechanical hypersensitivity. (**E**) Effect of the pan-Raf inhibitor TAK-632 on FL-induced mechanical hypersensitivity. Statistical analysis are performed using ANOVA two-way followed by Bonferroni’s test; ***P < 0.001 and ****P < 0.0001 compared to Vehicle-Saline (C) or Vehicle-FL group (D).All data are presented as mean ± SEM. Bsl: Baseline.

### FLT3-dependent ERK signaling pathways involved in the regulation of morphine analgesia

We previously reported that a functional interaction between MOR and FLT3 contributes to morphine-induced hyperalgesia and tolerance^8^. Several reports implicate ERK signaling in the development of analgesic tolerance and hyperalgesia^15^ and ERK is also critical in the transactivation of RTKs by G protein-coupled receptors (GPCRs) as well^24^. Hence, we investigated the activation of ERK signaling in HEK293T cells co-transfected with FLT3 and MOR after morphine and/or FLT3 application. As shown in Figure 5A, FL increased ERK phosphorylation, and this effect was significantly enhanced by co-application of morphine, whereas morphine alone did not significantly affect ERK phosphorylation. ERK phosphorylation was partially or totally reversed by the opioid receptor antagonist naloxone or the intracellular FLT3 inhibitor AC220, respectively confirming the specific implication of MOR and FLT3 in ERK activation. Three main canonical signaling pathways can be activated by FLT3 including PI3K/AKT/mTOR, JAK/STAT5, and MAPK/ERK1/2 pathway. To determine which signaling pathways were responsible for enhanced ERK activation after FL and morphine application, we assessed the effects of different pharmacological inhibitors on ERK activation. Our results showed that only the application of PD-98059 (10 µM) prevented ERK phosphorylation on HEK293T cells (Figure 5B).

**Figure 5.**
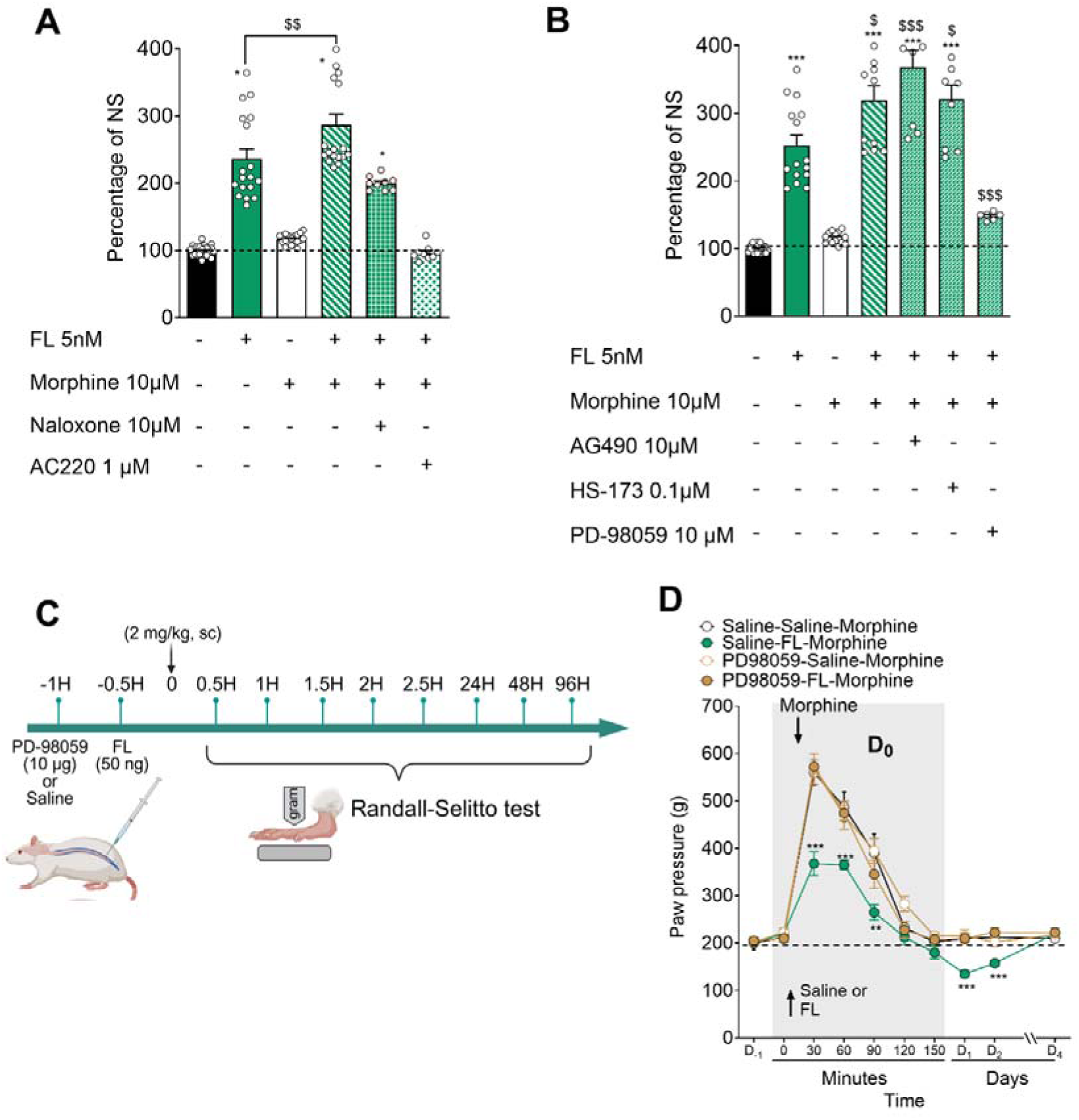
In vitro and in vivo assessment of the ERK signaling pathway in the functional interaction between FLT3 and MOR. (**A**) ERK1/2 activation in HEK293T cells expressing FLT3 and MOR following exposure to FL (5 nM), morphine (10 µM), or their combination. Cells were treated with the opioid receptor antagonist naloxone (10 µM) or the FLT3 inhibitor AC220 (1 µM). ERK1/2 activation is expressed as a percentage of the non-stimulated condition (NS), set at 100%. (**B**) ERK1/2 activation induced by combined FL (5 nM) and morphine (10 µM) exposure in the presence or absence of the JAK2 inhibitor AG490 (10 µM), the PI3K inhibitor HS-173 (0.1 µM), or the MEK inhibitor PD98059 (10 µM). Data are expressed as a percentage of the non-stimulated condition (NS), set at 100%. A-B; Statistical analysis was performed using ANOVA One-Way, followed by Bonferronni’s test. *, ** and ***P < 0.05, 0.01 and 0.001, respectively, versus NS condition; $, $$ and $$$ P < 0.05, 0.01 and 0.001, respectively, versus FL condition. (**C**) Experimental protocol used to assess the involvement of MEK-ERK signaling in the modulation of morphine-induced analgesia by FL. Rats received PD98059 (10 µg; i.t.) or saline 1 h before morphine (2 mg/kg; s.c.) administration and FL (50 ng; i.t.) or saline 30 min before morphine. Mechanical nociceptive thresholds were assessed using the Randall-Selitto test before treatment and from 30 min to 96 h after morphine administration. (**D**) Time course of paw-pressure thresholds in rats receiving morphine with or without FL pretreatment. Different groups received PD-98059 or vehicle injection. The grey area indicates the day of morphine administration (D□). Statistical analysis was performed using ANOVA Two-Way, followed by Bonferronni’s test. *, ** and ***P < 0.05, 0.01 and 0.001, respectively, versus Saline-Saline-Morphine group. Data are presented as mean ± SEM (n = 6 rats per group).

We then conducted *in vivo* experiments in naïve rats receiving intrathecal FL administration 30 minutes before morphine-2 mg/kg (Figure 5C) to determine whether ERK activation by FLT3 may alter morphine-induced analgesia. The administration of morphine increased the mechanical nociceptive threshold in vehicle-treated animals. As previously shown, morphine-induced increased paw pressure withdrawal is largely reduced by intrathecal FL pretreatment (Figure 5D) followed by sustained decreased mechanical threshold on D1 and D2. Whereas the intrathecal administration of PD-98059 (10 µg) did not change morphine-induced analgesia, it opposed to FL-induced decreased morphine analgesia and hyperalgesia (Figure 5D). Altogether, these data suggest synergistic activation of ERK pathway by FLT3 and MOR activation promoting decreased morphine analgesia and sustained mechanical pain hypersensitivity.

## Discussion

The present study identifies the Raf-MEK-ERK pathway as a major signaling mechanism downstream of FLT3 activation. Using complementary heterologous, primary neuronal, and *in vivo* approaches, we show that FL activates ERK through FLT3 and Raf, facilitates TRPV1-mediated calcium responses through MEK-ERK signaling, and induces ERK phosphorylation in the DRGs. We further demonstrate that pharmacological inhibition of Raf or MEK attenuates FL-induced mechanical hypersensitivity. Finally, our findings indicate that concomitant activation of FLT3 and MOR enhances ERK signaling and reduces morphine analgesic efficacy. These results extend our previous findings on the contribution of FLT3 to morphine tolerance and opioid-induced hyperalgesia by identifying Raf-MEK-ERK signaling as a potential mechanistic link between FLT3 activation and the modulation of MOR-mediated analgesia.

In FLT3-expressing HEK293T cells, both human and mouse FL induced a concentration-dependent increase in ERK phosphorylation, irrespective of the species origin of the receptor. These results indicate that ligand-dependent activation of the ERK pathway is conserved between mouse and human FLT3 and provide initial support for the potential translationnal relevance of the FL/FLT3 signaling mechanisms. They also validate the biological activity of the internally produced recombinant human FL used in our experiments. Activation of the Raf-MEK-ERK cascade downstream of FLT3 has primarily been characterized in hematopoietic and leukemic cells^21^. Our findings extend these observations by demonstrating that FLT3-dependent ERK activation can also occur in a non-hematopoietic cellular background. Moreover, the concentration-dependent inhibition produced by TAK-632 and the marked reduction in ERK phosphorylation produced by BDT001 support a signaling sequence in which FL binding activates FLT3 and subsequently recruits the Raf-MEK-ERK cascade.

Our results further establish a functional link between FLT3-dependent ERK signaling and the sensitization of TRPV1 in primary sensory neurons. We previously showed that FL enhances capsaicin-evoked intracellular calcium responses in DRG neurons through activation of neuronal FLT3^6^. The present data indicate that this effect requires, at least in part, MEK-dependent ERK activation. Importantly, PD-98059 did not modify the calcium response to capsaicin when administered alone, suggesting that MEK inhibition does not directly impair basal TRPV1 function or calcium influx under these experimental conditions. Instead, it selectively attenuated the potentiating effect of FL, supporting a role for the MEK-ERK pathway in FL-induced TRPV1 sensitization. These observations are consistent with previous studies showing that the MAPK pathway regulates TRPV1 function in sensory neurons. In particular, ERK activation contributes to TRPV1 sensitization by inflammatory mediators and neurotrophic factors^22,23^. More broadly, TRPV1 sensitization may involve channel phosphorylation, association with intracellular signaling complexes, and increased trafficking and insertion of the channel into the plasma membrane^25–27^. The present experiments do not distinguish between these possibilities. Nevertheless, the relatively rapid effect of FL favors a post-translational regulation of TRPV1 rather than a transcriptional increase in channel expression. Further studies assessing TRPV1 phosphorylation and surface expression will be required to define the molecular mechanism through which FLT3-dependent ERK activation enhances channel function. The relevance of this signaling pathway *in vivo* is supported by the rapid increase in ERK phosphorylation observed in the DRGs following intrathecal FL administration. The temporal profile of this response detected within 30 min and maintained at 60 min, is compatible with direct activation of FLT3-dependent intracellular signaling in primary sensory neurons. Consistent with our previous studies^6,7,9^, intrathecal FL also produced robust mechanical hypersensitivity. This behavioral effect was attenuated by both TAK-632 and PD-98059, indicating that sequential activation of Raf and MEK-ERK contributes to the pronociceptive effects of FL *in vivo*. Together with the calcium-imaging data, these findings support a model in which FLT3 activation increases the responsiveness of nociceptive neurons through Raf-MEK-ERK-dependent modulation of excitability and ion-channel function.

Our findings also provide new insight into the interaction between FLT3 and MOR signaling. We previously demonstrated that FLT3 functionally interacts with MOR and contributes to the development of morphine-induced hyperalgesia and analgesic tolerance^8^. Here, morphine enhanced FL-induced ERK phosphorylation in HEK293T cells co-expressing FLT3 and MOR. This enhanced response was partially reduced by naloxone and totally abolished by the intracellular FLT3 inhibitor AC220. These pharmacological data suggest that MOR activation facilitates FLT3-dependent ERK signaling. Several mechanisms could account for this functional interaction. GPCRs can regulate receptor tyrosine kinase signaling through transactivation, receptor-associated signaling complexes, or convergence of their downstream pathways at the level of Ras, Raf, or MEK^28–30^. Conversely, FLT3 activation may modify MOR signaling, trafficking, or desensitization. The complete blockade of ERK phosphorylation by AC220 suggests that, under the present experimental conditions, the increase produced by combined MOR and FLT3 activation is strongly dependent on FLT3 kinase activity. However, these results do not yet demonstrate direct MOR-FLT3 transactivation or physical receptor interaction. Additional experiments will be necessary to distinguish direct transactivation from convergence of downstream signaling pathways between MOR and FLT3.

The functional relevance of ERK activation to opioid responses is supported by the observation that PD-98059 preserved morphine analgesia and prevented the mechanical hypersensitivity induced by the combination of morphine and FL. These findings are consistent with the established involvement of ERK signaling in opioid receptor plasticity, analgesic tolerance, and opioid-induced hyperalgesia^15–17^. In line with our results, Jin et al. recently identified sensory-neuron BRAF as a critical mediator of opioid-induced hyperalgesia and tolerance. They showed that morphine promotes the translocation of monomeric BRAF from the DRG to nociceptor central terminals, where it increases MEK-ERK activation and enhances the phosphorylation, synaptic trafficking, and activity of presynaptic NMDA receptors. Pharmacological inhibition of BRAF or MEK, as well as sensory-neuron-specific deletion of *Braf*, enhanced morphine analgesia while attenuating opioid-induced hyperalgesia and tolerance^31^. Because BRAF is a canonical downstream effector of tyrosine kinases, FLT3 may provide an upstream signal that regulates BRAF-MEK-ERK activity in sensory neurons, although a direct functional link between FLT3 and BRAF remains to be established. Accordingly, inhibition of FLT3 or its downstream MAPK pathway may preserve opioid analgesia while limiting the development of maladaptive responses to repeated opioid exposure^8^. Finally, the *in vivo* experiments were conducted in male mice, and potential sex-dependent differences remain to be investigated.

In conclusion, this study identifies the Raf-MEK-ERK pathway as a functional signaling axis downstream of FLT3 in sensory neurons. Activation of this pathway contributes to FL-induced TRPV1 sensitization and mechanical hypersensitivity and appears to mediate the detrimental influence of FLT3 signaling on morphine analgesia. These findings extend the role of neuronal FLT3 beyond the direct regulation of nociceptor excitability and position FLT3-dependent ERK signaling as a potential mechanistic link between sensory-neuron sensitization, pain hypersensitivity, and opioid-induced plasticity. More broadly, characterization of the intracellular pathways engaged by neuronal FLT3 provides a better understanding of its pronociceptive effects and of the mechanisms through which it may modulate morphine efficacy.

## Acknowledgements

We thank the different technical platforms of Institut des Neurosciences de Montpellier (INM), especially the animal care facility and the functional exploration platform RAM Neuro supervised by Denis Greuet. This work was supported by INM, INSERM, Montpellier University and grants from: La Région Languedoc-Roussillon, the SATT AxLR, National Research Agency (grant ANR-19-CE16-0008 to C.R.), Biodol Therapeutics. We thank Professor Rachel Cerdan (Laboratory of Pathogen Host Interactions INSERM-CNRS, University of Montpellier, France) for her critical comments and expert advice on the structural comparison of human and mouse FLT3.

